# The Impact of Graph Construction Scheme and Community Detection Algorithm on the Repeatability of Community and Hub Identification in Structural Brain Networks

**DOI:** 10.1101/2020.05.07.082271

**Authors:** Stavros I. Dimitriadis, Eirini Messaritaki, Derek K. Jones

## Abstract

A critical question in network neuroscience is how nodes cluster together to form communities, to form the mesoscale organization of the brain. Various algorithms have been proposed for identifying such communities, each identifying different communities within the same network. Here, (using test-retest data from the Human Connectome Project), the repeatability of 33 community detection algorithms, each paired with 7 different graph construction schemes was assessed.

Repeatability of community partition depended heavily on both the community detection algorithm and graph construction scheme. Hard community detection algorithms (in which each node is assigned to only one community) outperformed soft ones (in which each node can be belong to more than one community). The highest repeatability was observed for the fast multi-scale community detection algorithm paired with a graph construction scheme that combines 9 white matter metrics. This pair also gave the highest similarity between representative group community affiliation and individual community affiliation. Connector hubs had higher repeatability than provincial hubs. Our results provide a workflow for repeatable identification of structural brain networks communities, based on optimal pairing of community detection algorithm and graph construction scheme.

## 1. Introduction

The human brain can be modelled as a network (Bassett and Sporns, 2017) and summarised as a graph. In structural networks, the nodes of the graph are small volumes of tissue which are interconnected via white matter tracts (edges). Graph theory can provide novel insights into healthy human brain function (Bassett et al., 2011; Braun et al., 2015) and its alteration in various diseases (Braun et al., 2016, Baker et al. (2015); Collin et al. (2016); Drakesmith et al. (2015); Aerts et al. (2016); Nelson et al. (2017); Vidaurre et al. (2018); Imms et al. (2019)).

An open question in network neuroscience is how neural units cluster together to form inter-connected groups and provide the coordinated brain activity that gives rise to action, perception and behaviour (Bassett and Mattar, 2017; Pessoa et al., 2018). Modularity is a quintessential concept in network neuroscience, wherein neural units are densely connected to one another, forming clusters or *modules* (Meunier et al., 2010). This is an efficient architecture allowing a complex network to integrate information locally, while maintaining its adaptability to any external stimulus. Networks in nature often show hierarchical, modular organization (Blondel et al., 2008; Fortunato 2010; Fortunato and Castellano 2012; A. Lancichinetti and Fortunato 2009a,b; Newman,2004 2012; Meunier et al., 2009). In the brain, such hierarchical modularity could support segregated neuronal interactions and their integration at the global level. Networks with such structure (Fortunato, 2010) are more complex than those with random structure (Sporns et al., 2000), and have been well demonstrated in functional brain networks (Sporns and Betzel, 2016).

Commonly studied graph theoretical metrics, such as global and local efficiency, clustering coefficient, shortest path length, and small-worldness (Rubinov and Sporns 2010), provide useful information related to the global and local properties of the network, but to investigate *mesoscale* network organization community (or modularity) detection techniques are more appropriate (Meunier et al. 2010; Giusti et al., 2016). Following community partition, a frequently used methodology to identify the community structure uses two modular network metrics (Guimera et al., 2005; van den Heuvel and Sporns, 2013). The participation coefficient, P_i_, of a node quantifies the distribution of its links among the modules of the network, while the within-module z-score, z_i,_ measures the connectedness of a node to other nodes in the module. These metrics in turn can be used to classify hubs as either provincial or connector hubs (Guimera and Amaral, 2005). Please refer to Appendix 1 for more detail on the computation of P_i_ and z_i_ and how this classification is made.

Complex network analysis of multimodal structural and functional brain connectivity has identified a subset of brain areas that play a key role for an efficient neural signalling and communication (van den Heuvel and Sporns, 2013). These brain areas, called hubs, support dynamic functional coupling within and between functional subnetworks. In empirical structural brain networks, the term ‘rich-club’ characterises brain areas/nodes with high degree that are more densely interconnected between each other compared to the rest of the network.

Various methodologies for structural network generation have been proposed, involving tractography with different algorithms and assigning edge weights using different diffusion MRI-based metrics. The resulting graphs are quite different from each other and have different levels of robustness and repeatability (Smith et al., 2015; Owen et al., 2013, 2013; Zhong et al., 2015; Dimitriadis et al., 2017b; Yuan et al., 2019). We recently explored the repeatability of structural brain graphs, their edge weights and graph-theoretical metrics, for twenty-one different edge-weighting schemes (Messaritaki et al., 2019a). We demonstrated that integrating several metrics as edge weights is very good at capturing differences between populations, and is interesting from the perspective of developing biomarkers (Dimitriadis et al 2017, Clarke et al 2020).

We constructed structural brain networks from a set of test-retest diffusion MRI scan data from the Human Connectome project using the b=2000◻s/mm^2^ data and the seven most reproducible graph-construction schemes as derived from our previous study on the same data (Messaritaki et al., 2019b). We then applied thirty-three community detection algorithms. The ‘hard’ algorithms assign every node to only one community, while the ‘soft’ algorithms can assign a node to multiple communities. For every pair of community detection algorithm and graph construction scheme, we estimated the reproducibility of nodal P_i_ and z_i_ and of provincial and connector hubs, based on both modular network metrics. Our aim was to identify the combination of graph construction scheme and community detection algorithm with the highest agreement of individual communities between the two repeat scan sessions.

The quality criterion for the estimated community partitions was also important in our study. To this end, we compared the quality index of the community partitions estimated over the original graphs with quality indexes of the community partitions computed over surrogate null versions of the original graph (Guimera et al., 2004). We previously reported a statistical procedure for performing condition and group comparisons in terms of brain communities (Dimitriadis et al., 2012). Here, we applied a similar approach to assess between-scan pairwise community similarity for every pair of graph construction scheme and community detection algorithm. We adopted a proper community partition distance metric, the Normalized Mutual Information (NMI) (Lancichinetti and Fortunato,2009a ; Alexander-Bloch et al. 2012). Finally, we derived a consensus cluster across participants and repeat scans (Dong et al., 2014; Ozdemir et al., 2015). The agreement of consensus cluster with individual communities adopting NMI was also used as an objective criterion of the optimal combination of graph construction scheme and community detection algorithm.

We note that the analysis presented here does not aim to assess how well these structural networks represent the organisation of the human brain. The accuracy of these networks and of the metrics used as edge-weights in representing the functional organisation of the brain has been validated in recent work by Messaritaki et al. (2020). Additionally, the metrics used as edge-weights are routinely used in network analyses in the literature (for example Taylor et al., 2015; Nigro et al., 2016; Caeyenberghs et al., 2016). Our analysis does, however, address one aspect of the accuracy of the partition of the structural connectome. If a partition of the structural connectome is not repeatable in the absence of changes resulting from maturation or intervention, then that partition is not an accurate representation of the modular organisation of the structural connectome. Only partitions that are repeatable can convey reliable information about the structural organisation of the human brain. In other words, even though the repeatability of a partition is not a sufficient condition for it to be representative of the brain’s structural organisation, it is a necessary one.

The rest of this manuscript is organised as follows: Section 2 describes the graph-construction schemes, community detection algorithms, community partition similarity, the methodology for detecting connector and provincial hubs and their repeatability. Section 3 reports our results in terms of repeatable community partitions across the 2D space of graph-construction schemes / community detection algorithms, the repeatability of nodal P_i_/ z_i_ and the detection of connector/provincial hubs. The Discussion summarises the main outcome of our study explaining its advantages, limitations, and suggestions for future directions.

## 2. Methods

All analyses were performed using MATLAB (2019a; The Mathworks, Inc, Massachussets, United States).

### 2.1. Data

We analysed the test-retest MRI and diffusion-MRI dataset from the multimodal neuroimaging database of the Human Connectome Project (HCP) (Sotiropoulos et al., 2013b; Glasser et al., 2013 ; van Essen et al., 2013). We used the data from the 37 participants for whom there were 90 gradient directions for each *b*-value. The participants on this test-retest dataset were scanned twice with the between-scan time interval ranging between 1.5 and 11 months. The age-range of the participants was 22–41 years. The test-retest time interval is shorter than the expected time over which maturation-induced structural changes can be measured with diffusion MRI (dMRI).

The diffusion-weighted images (DWIs) had a resolution of (1.25×1.25×1.25) mm^3^ and were acquired for three different diffusion weightings (*b*-values: 1000◻s/mm^2^, 2000◻s/mm^2^ and 3000◻s/mm^2^) across 90 gradient directions for each *b*-value. The HCP acquisition details and pre-processing are described in Sotiropoulos et al. (2013a,b), Feinberg et al. (2010), Moeller et al. (2010), Setsompop et al. (2012), Xu et al. (2012), Glasser et al. (2013). Specifically, the diffusion images were corrected for EPI distortions, eddy-current distortions, participant movement and gradient nonlinearities. The diffusion data were also registered to the structural data. We performed the following analyses using the b=2000◻s/mm^2^ data.

### 2.2. Tractography

We performed whole-brain tractography using ExploreDTI-4.8.6 (Leemans et al., 2009), estimating the fiber orientation distribution function (fODF) using constrained spherical deconvolution (CSD) (Tournier et al., 2004). Tracking was initiated on a 2×2×2 mm grid, with a 1mm step size, angular threshold of 30° and fiber length range of 50−500 mm.

### 2.3. Graph generation

#### 2.3.1. Node definition

The Automated Anatomical Labeling (AAL) atlas (Tzourio-Mazoyer et al., 2002) was used to define 90 cortical and subcortical areas (45 areas per hemisphere) as nodes of the structural brain graphs. Structural brain networks (SBN) were generated for each participant using ExploreDTI-4.8.6 (Leemans et al., 2009).

#### 2.3.2. Edge weights

Edges were weighted using the seven most reproducible graph-construction schemes identified previously with the same dataset (Messaritaki et al., 2019b), and which were based on different combinations of the nine metrics listed in Table 1 (see Section 2.3.4). Each graph was normalised to have a maximum edge weight of 1, while the elements in the main diagonal were set to zero (see Figure 1).

**Table 1.** Metrics used in connectivity matrices.

| **Metric** | **Abbreviation** |
| --- | --- |
| Fractional anisotropy | FA |
| Mean diffusivity | MD |
| Radial diffusivity | RD |
| Number of streamlines | NS |
| Percentage of streamlines | PS |
| Streamline density | SLD |
| Tract volume | TV |
| Tract length | TL |
| Euclidean distance between nodes | ED |

**Figure 1.**
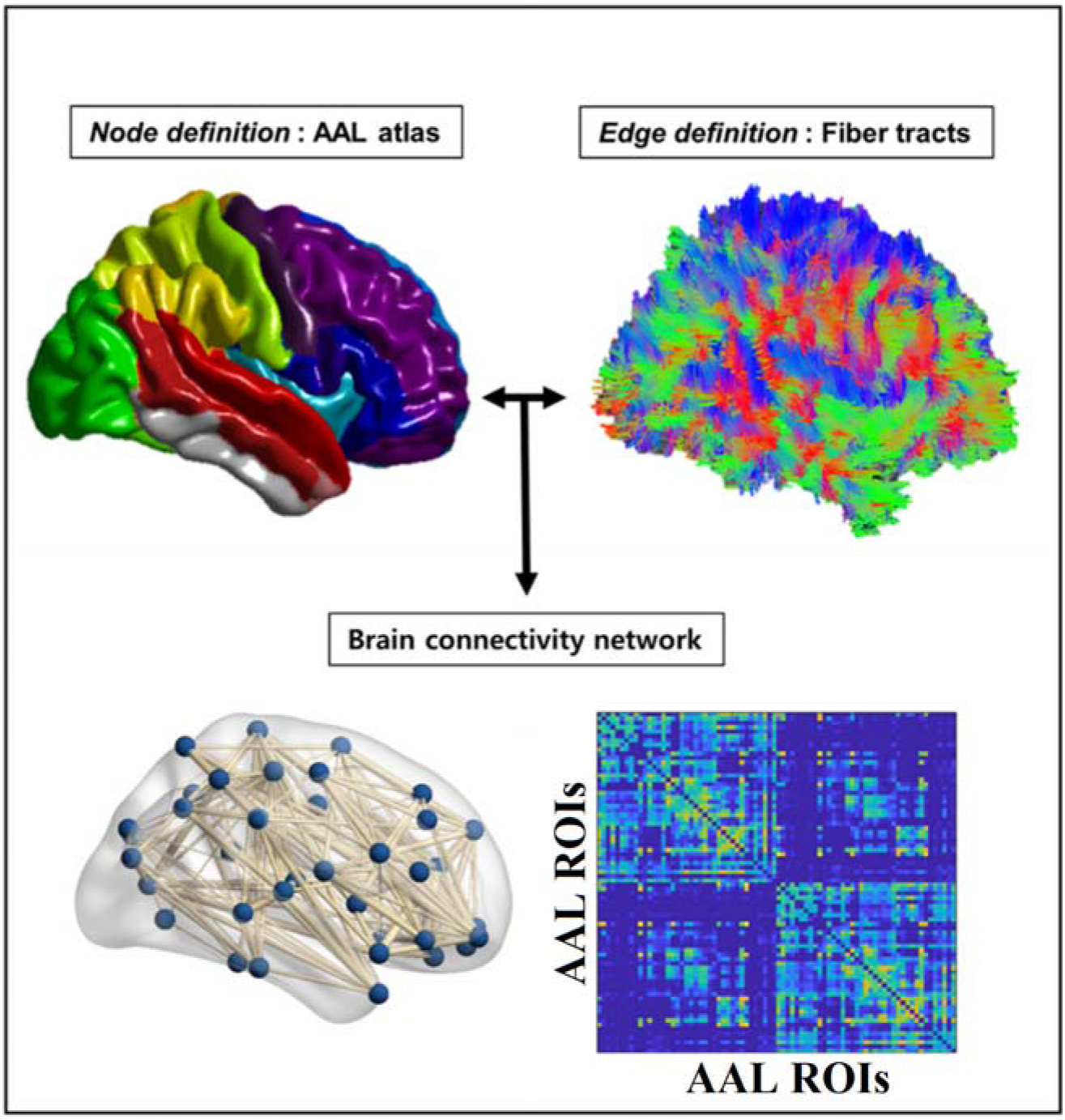
Flowchart of the construction of a structural brain network based on tractography and diffusion metrics (see Table 1).

#### 2.3.3 Integrated Edge-Weights

Combining multiple metrics into an *integrated* edge weight is supported by the fact that each metric conveys information about different tissue properties, while at the same time topological properties of SBNs are affected by more than one metric. Here, using the data-driven algorithm described in our previous work (Dimitriadis et al., 2017b,c) the nine metrics in Table 1 were used to form an integrated SBN for each participant and scan session.

The algorithm down-weights the more similar metrics and up-weights the most dissimilar metrics to enhance the integration of complementary topological information across the nine metrics. An orthogonal-minimal-spanning-tree (OMST) algorithm was then applied to the resulting networks, selecting the edges that preserve connectivity between nodes, while guaranteeing that the overall network efficiency is maximised. More details on the OMST algorithm and its implementation can be found in our previous work (Dimitriadis et al., 2017a,b,c; Messaritaki et al., 2019) and the related code is freely available at https://github.com/stdimitr/multi-group-analysis-OMST-GDD.

#### 2.3.4 Graph Construction Schemes

Seven graph construction schemes were used in this study, summarised in Table 2 and falling broadly into two categories. We briefly explain their construction methodologies here.

**Table 2.** Summary of the graph-construction schemes.

| Abbreviation | Initial Edge Weights | Topology | Final Edge Weights | Symbol |
| --- | --- | --- | --- | --- |
| NS – OMST | NS | OMST | Unchanged | A |
| NS + FA OMST | lin. comb. of NS and FA | OMST | Unchanged | B |
| 9-m OMST | lin. comb. of all 9 metrics in Table 1 | OMST | Unchanged | C |
| NS-thr | NS | keep highest-NS edges | Unchanged | D |
| NS-t/FA-w | NS | keep highest-NS edges | re-weight with FA | E |
| NS-t/MD-w | NS | keep highest-NS edges | re-weight with MD | F |
| FA-t/NS-w | FA | keep highest-FA edges | re-weight with NS | G |

The first category includes graphs constructed via the data-driven algorithm (Dimitriadis et al., 2017b, c). A) NS-OMST: apply the OMST filtering algorithm (Dimitriadis et al., 2017b,c) to the NS-weighted matrix. B) NS+FA - OMST: Integrate the NS-weighted and FA-weighted matrices with the data-driven algorithm. C) 9m-OMST: Integrate all nine diffusion metrics (as originally reported in Dimitriadis et al. 2017b, see Table 2).

The second category includes SBNs with edges weighted by the NS or the FA and applying a threshold to remove edges with the lowest weights. The threshold was determined by imposing the constraint that the graphs exhibit the same sparsity as the OMST graphs that exhibited the highest reproducibility (Messaritaki et al., 2019b). Once the topology of each of those graphs was specified, the weights of the edges were either kept as they were or re-weighted with one of the remaining two metrics. These graphs are as follows (see Table 2). D) NS-thr: Keep the highest-NS edges. E) NS-t/FA-w: Threshold to keep the highest-NS edges, then reweight those edges with their FA. F) NS-t/MD-w: Keep the highest-NS edges, then reweight those edges with their MD. G) FA-t/NS-w: Keep the highest-FA edges, then reweight those edges with their NS.

As we have shown previously, these seven schemes exhibit different values of similarity between them, from 0.99 to 0.42 (Messaritaki et al. (2019b), Table 3), motivating their inclusion in a study on the repeatability of community detection.

**Table 3.** Group-averaged similarity of individual community partitions with consensus community partition. Similarities are expressed in NMI scale. We assigned with bold the top ranked values.

|  | NS-OMST | NS+FA<br>OMST | 9-m<br>OMST | NS-thr | NS-t/FA-w | NS-t/MD-w | FA-t/NS-w |
| --- | --- | --- | --- | --- | --- | --- | --- |
| mscd_afg | 0.64 | 0.57 | **0.70** | 0.61 | 0.61 | 0.61 | 0.39 |
| mscd_rb | 0.53 | 0.39 | 0.39 | 0.32 | 0.31 | 0.30 | 0.18 |
| mscd_rn | 0 | 0 | 0.36 | 0 | 0 | 0 | 0 |
| mscd_so | **0.70** | 0.65 | **0.75** | **0.68** | **0.68** | **0.68** | 0.64 |

#### 2.3.5 Community detection algorithms

Communities or modules are defined as subgroups of nodes that are more interconnected with each other compared to the rest of the network (Newman and Girvan 2004; Radicchi et al. 2004). In the present study, we compared thirty-three different community detection algorithms, comprising twenty-six with hard clustering and seven with soft clustering. (see Figure 2). In hard clustering, community membership can be represented as a vector that encapsulates the assignments of every brain area to every detected graph cluster (community). In our case, clustering has a dimension of 1 x 90, equalling the number of brain regions in the AAL parcellation. In soft clustering, the outcome is a matrix that encapsulates how many soft clusters a given node (brain area) belongs to. A more detailed description of the adopted community detection algorithms is provided in Appendix 2 & Supplementary Material.

**Figure 2.**
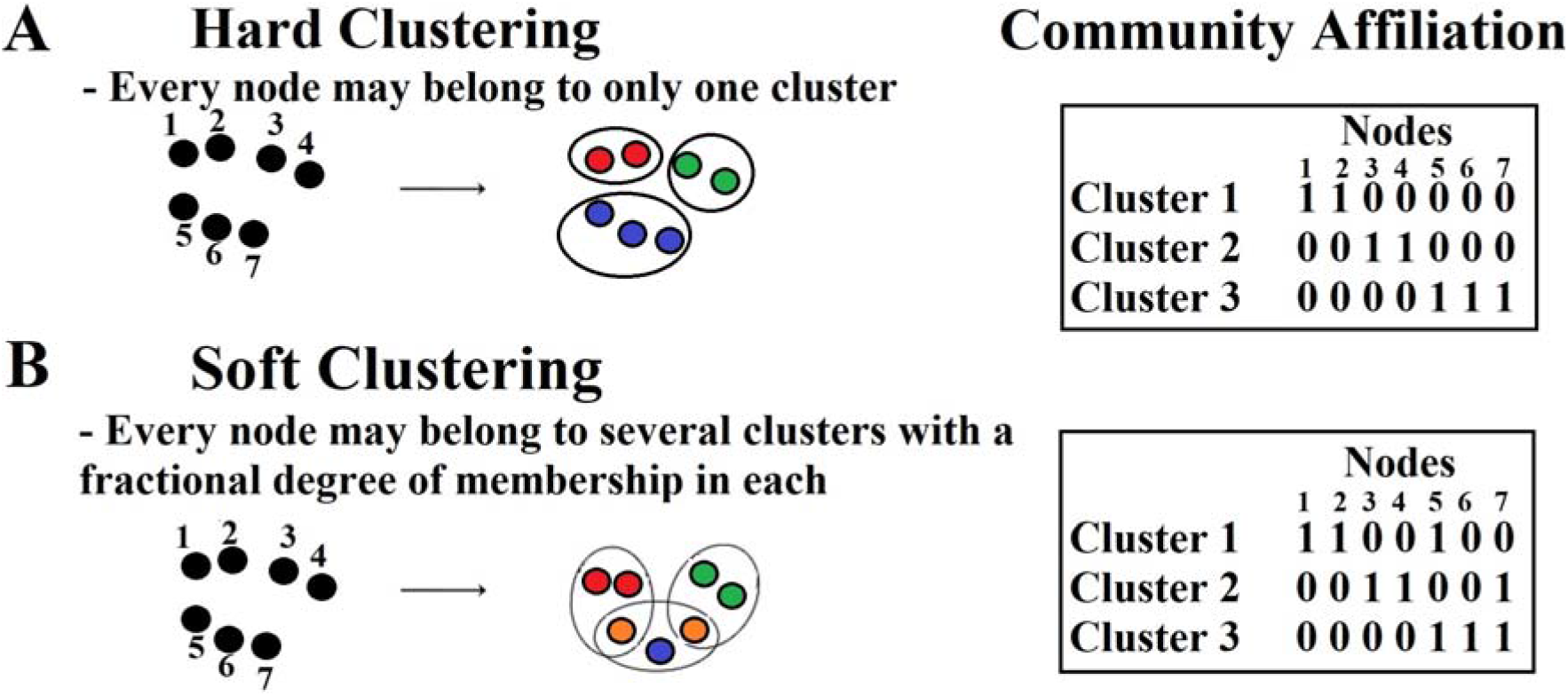
An example of hard and soft clustering in a toy example containing 7 nodes. A. Hard clustering: A node can only belong to one cluster. The table shows the community assignment to every node. B. Soft clustering: Five out of seven nodes are clustered in a single cluster/community {nodes 1,2,3,4,6} while nodes {5 and 7} belong to two communities: node 5 belongs to communities 1 and 3 while node 7 belongs to communities 2 and 3. The table shows the community assignment to every node.

In the present study, we considered for the very first time in brain network neuroimaging a large number of community detection algorithms. We adopted thirty-three graph partition algorithms further divided into twenty-six hard clustering algorithms and seven soft clustering algorithms.

#### 2.3.6 Permutation Test on Quality Modular Indexes

For every participant, scan and graph construction method, we produced 1000 surrogate null graph models by randomizing the weighted connections while preserving both the degree and strength of every node and the overall connectedness of the network (Rubinov and Sporns,2010).

All of the hard clustering algorithms (no.s 1-26) involved a Q quality index for the communities detected. For further details of Q quality indexes see Le Martelot and Hankin, (2011,2012 a,b).

For the soft clustering algorithms (no. 27-33), we estimated the Normalized Mutual Information (NMI; see Appendix 3) between the original community affiliation and the surrogate null communities produced via the application of every algorithm to the surrogate graph model.

#### 2.3.7 Between-Scan Community Detection Agreement

We quantified the graph-partition distance with the normalized mutual information (NMI; see Appendix 3).

#### 2.3.8 Consensus Clustering

A consensus matrix was constructed for every pair of {graph construction scheme – community detection algorithm} that showed high test-retest reproducibility across the cohort (group-averaged NMI > 0.9). We quantified how many times two nodes across the 74 SBNs (37 participants x 2 scans) were classified in the same community and this entry, t_i,j,_ was assigned to the relevant pair of nodes. The consensus matrix has the same dimensions as the original SBN (90 x 90 in our case) with entries assuming integer values between 0 and 74 {37 participants x 2 scans}, which were then transformed to denote the probability of occurrence of a pair of nodes (brain areas) being classified as belonging to the same community across the cohort and scan sessions. We converted the consensus matrix into a probability one by dividing each entry by 74.

In order to get a consensus or group representative community per graph-construction scheme and community detection algorithm, consensus matrices should be iteratively thresholded and clustered with a community detection algorithm (Lancichinetti and Fortunato, 2012). This algorithm uses an absolute arbitrary threshold to eliminate weak connections and iteratively apply a graph partition technique. Instead of an arbitrary filtering scheme, we adopted our OMST algorithm (Dimitriadis et al., 2017a,b,c) to topologically-filter the consensus matrix in a data-driven way. We then extracted the consensus – group representative community by applying the community detection algorithms across the graph construction schemes (Newman,2006). See Figure 3.E for an example of a consensus matrix.

**Figure 3.**
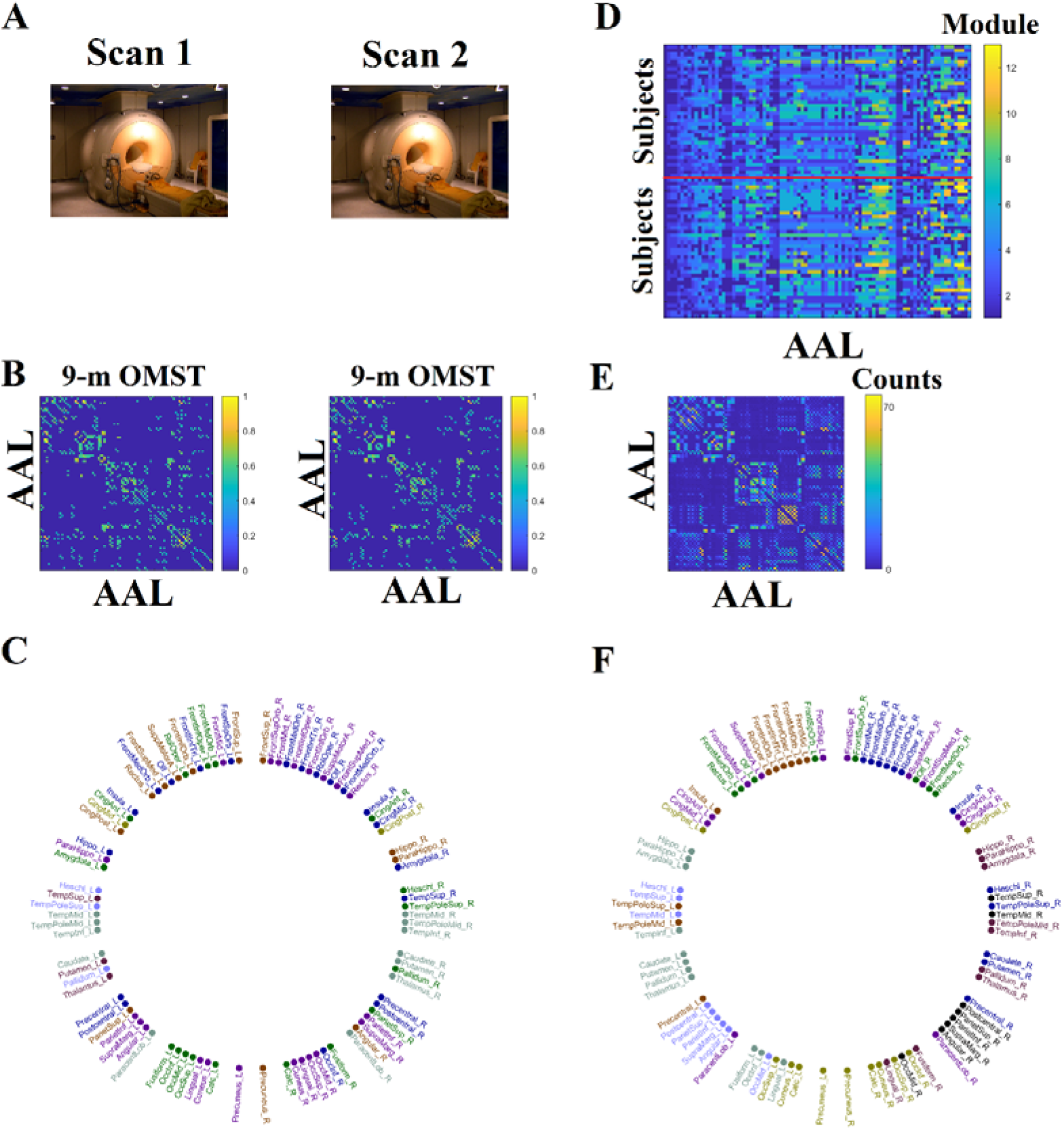
Outline of the presented methodology. **The demonstration based on 9-m OMST graph-construction scheme and gso-discrete mode community detection algorithm.** A. Repeat – Scan Sessions B. Structural brain networks from participant 1 from both sessions using 9-m OMST graph-construction scheme C. Individual community affiliation of participant 1 from scan session 1. Each colour represents one community. D. Vectorised community affiliations of the whole cohort from scan sessions 1 and 2 separated with a red line. Every module is coded with a different colour. E. Consensus matrix is built over group community affiliations across both sessions as presented in D. Weights in the consensus matrix refer to the total number of times two brain areas are grouped together across the cohort and scan sessions with the maximum value being (number of participants) × (scan sessions) = 74. F. Representative community affiliation after graph partitioning the consensus matrix presented in E. Each community is encoded to a different colour. Similarity NMI distance has been estimated between representative community affiliation presented in F and individual community affiliations presented in C.

#### 2.3.9 Agreement of consensus representative community with individual community structures

An important criterion of our analysis is the high similarity between the consensus clustering and individual clustering for every graph construction scheme that showed high group-averaged community similarity (NMI > 0.9). To this end, we estimated this community similarity for every case.

Figure 3 illustrates the various steps of the analysis.

#### 2.3.10 Evaluating the Combined Graph Construction Schemes - Community Detection Algorithms

As mentioned previously, we first identified the combinations of graph construction schemes and community detection algorithms with higher group-averaged between-scan community affiliation agreement (NMI > 0.9) with a p < 0.05 based on the bootstrapping procedure. We then adopted a criterion of highest community similarity between the consensus clustering with individual community affiliation (clustering). It is important that consensus clustering expresses the inter-subject variability and acts as a vector median for the whole group (Dimitriadis et al., 2012). The final ranking of pairs of graph construction schemes and community detection algorithms was based on: 1) high between-scan group-averaged community similarity with a p < 0.05 (bootstrapping), 2) Q quality index with p < 0.05 based on surrogate null brain models and 3) high community similarity between consensus clustering and individual community affiliations (clusterings) assessed via a two-way ANOVA (see section 2.7).

### 2.4 Modular Driven Structural Brain Hub Detection

Here, we applied the aforementioned methodology solely on the graph construction schemes and community detection algorithm that fulfil the evaluation criteria of section 2.3.10.

### 2.5 Reliability of Nodal Participation Coefficient P_i_ and Within-Module Z-Score z_i_

We also explored the intra-class correlation coefficient (ICC) of nodal participation coefficient *P_i_* and within-module z-score *z_i_*. As a main outcome of this hub detection approach, we quantified the consistency of connector/provincial hub detection first within participant between scans, and secondly across the cohort.

### 2.6 Assessing a Reproducible Structural Core of the Human Brain

We detected structural hubs for every participant, scan session and graph construction scheme by applying an absolute threshold to the participation coefficient and within-module z-score (Guimera and Amaral,2005; Hagmann et al., 2007,2008); van den Heuvel and Sporns,2013). We estimated an agreement index that quantifies the percentage of connector/provincial hubs that were detected in both scans. This agreement index is defined as:

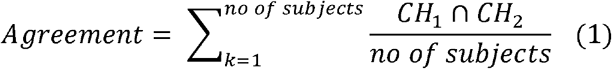

where CH_1,2_ are two vectors of size 1×90 with ones in positions where a brain area is detected as connector or provincial hub. This agreement index is normalized by the total number of participants and takes the absolute value of 1 when a node/ROI is detected as either connector or provincial hub across all participants and in both scans.

### 2.7 Statistical Analysis

Firstly, we determined pairs of graph construction schemes and community detection algorithms that fulfilled the evaluation criteria presented in 2.3.10. The first two criteria were evaluated via bootstrapping and surrogate null models, while for the third, based on the group-averaged similarity of individual community partitions with consensus community partition, a two-way ANOVA was adopted (p < 0.05).

The detection of reproducible brain structural hubs employing community-based hub detection network metrics require reproducible communities. For that reason, we followed hub detection analysis over the best pairs of graph construction schemes and community detection algorithms. Then, we adopted a two-way ANOVA (p < 0.05) to detect the best pair of graph construction scheme and community detection algorithms over hub detection analysis, using as input the ICC for the participation index and the within-module z-score across nodes and the agreement-index.

## 3. Results

### 3.1 Quality of the Detected Communities

Q original values were transformed into p-values by comparing them with the 1000 surrogate (permuted) Q values (see sup.material). P-values ranged between 0.013 and 0.021 across participants, scans and graph construction schemes. For the soft clustering algorithms (no. 27-33) that do not include a Q quality index, we measured the NMI between the original community affiliation and the 1000 surrogate-based communities. The group-averaged NMI ranged between 0.08 and 0.12 which supported the quality of the derived soft communities with the whole set of soft clustering algorithms. These findings support the quality of the extracted graph partitions and allow us to include all the participants, scans, graph construction schemes and community detection algorithms in our analysis.

### 3.2 Group-Averaged Between-Scan Agreement of Communities Affiliations

Figure 4 shows group-averaged between-scan agreement of community affiliations across graph constructions schemes and community detection schemes. Based on the highest group-averaged NMI values and the detected p-values (p = 0.0001) derived from the permutation test, we detected the following four community detection algorithms across the seven graph construction schemes as having the highest between-scan agreement:

1. (<u>mscd_afg</u>): Fast multi-scale community detection algorithm using the criterion from Arenas et al. (2008)
2. (<u>mscd_rb</u>): Fast multi-scale community detection algorithm using the criterion from Reichardt & Bornholdt, (2006)
3. (<u>mscd_rn</u>): Fast multi-scale community detection algorithm using the criterion from Ronhovde & Nussinov (2009)
4. (<u>mscd_so</u>): Fast multi-scale community detection algorithm using stability optimisation (Le Martelot and Hankin, 2012).

**Figure 4.**
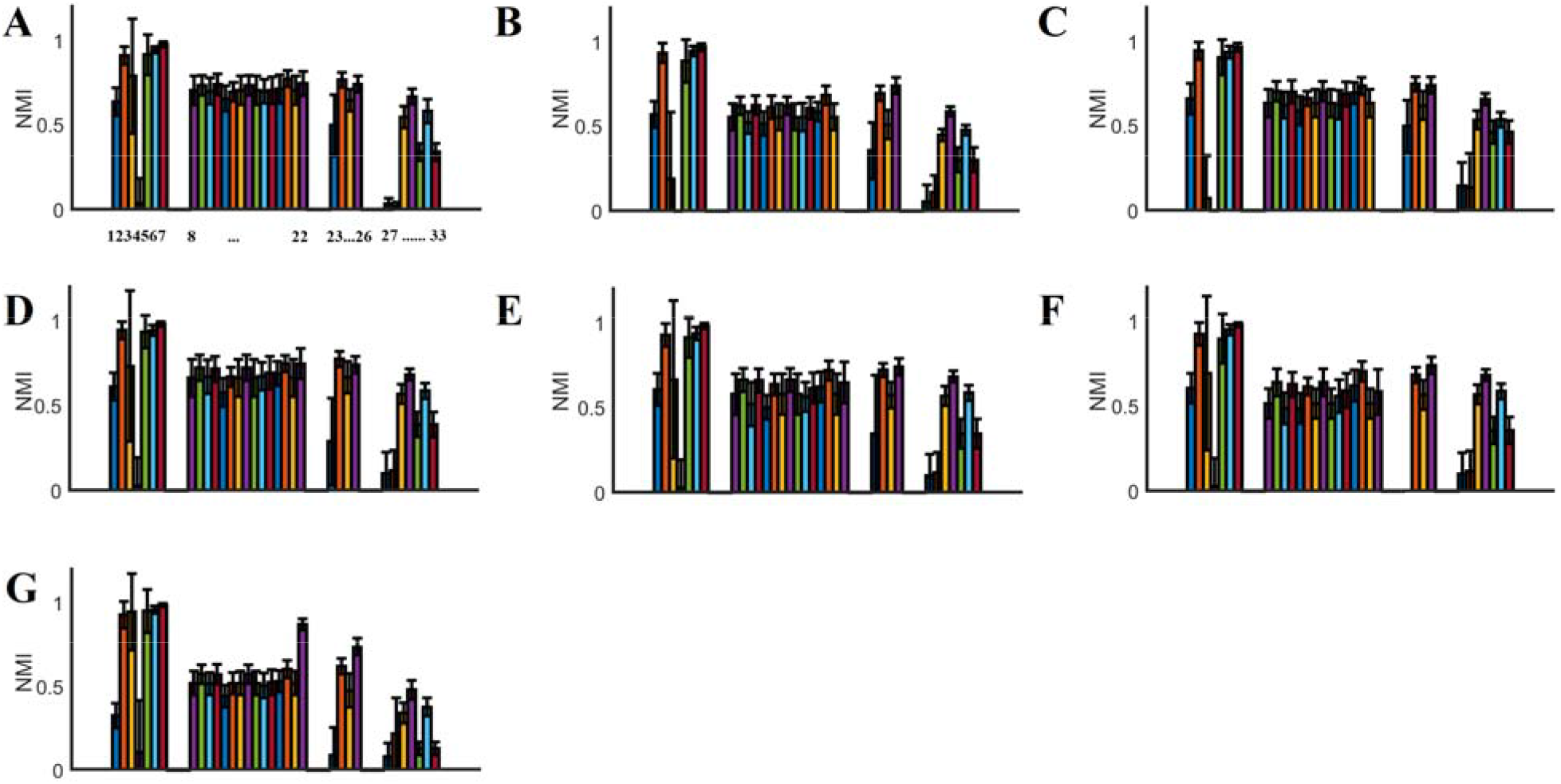
Between-Scan Agreement of Communities Affiliations across graph construction schemes and community detection algorithms. Every subplot refers to one of the 7 graph-construction schemes. The bars define the group-averaged between-scan agreement of community affiliations. Numbers below the plot in A refer to the number list of community detection algorithms represented in section 2.3.3. Community detection algorithms with the highest agreement between the two scans (NM1 > 0.9) were: mscd_afg,mscd_rb, mscd_rn and mscd_so. For the abbreviations of the graph-construction schemes please see Table 2.

### 3.3 Similarity of Individual Community Partitions with Consensus Community Partition

The highest similarity between individual community partitions and consensus community partition was detected for the combination of 9-m-OMST graph construction scheme and mscd_so community detection algorithm. The second highest similarity was detected for mscd_afg and 9-m OMST. MSCD with *rb* and *rn* criterions failed to produce an acceptable community similarity between a group representative community estimated via consensus clustering and the individual community affiliations (see Table 3). Two-way ANOVA reported an effect of group-averaged similarity of individual community partitions with consensus community partition across the four community detection algorithms (df =3, F = 54.14, p = 0.783 x 10^−9^, corrected for multiple comparison) but there was no effect of the graph construction or their interaction. Based on the mean group-averaged similarity across the seven graph construction schemes, we ranked the four community-detection algorithms. The Mscd_so community detection algorithm produced the highest group-averaged similarity of individual community partitions with consensus community partition compared to the four community detection algorithms.

### 3.4 Evaluation of the Best Combination of Graph Construction Scheme and Community Detection Algorithm

Based on the evaluation criteria presented in section 2.4 for the final ranking of the pairs of graph construction schemes and community detection algorithms, and taking into account the individual scores, we ranked as the best pair the combination of mscd_so with 9-m OMST. This pair showed a high between-scan group-averaged community similarity estimated with NMI and evaluated via bootstrapping (p = 0.0001). The Q quality index has a value of 0.67 with p = 0.001. The Mscd_so community detection algorithm also produced the highest group-averaged similarity of individual community partitions with consensus community partition compared to the four community detection algorithms supported by two-way ANOVA.

Figure 5 illustrates the topology of the nine communities across the 90 AAL brain areas based on the combination of mscd_so with 9-m OMST. Importantly, modules number 1, 7 and 9 group together brain areas located within the left hemisphere, modules number 2, 6 and 8 group together brain areas located within the right hemisphere, while modules 3, 4 and 5 involve areas from both hemispheres. Specifically, they involve bilateral ROIs from the fronto-parietal, cingulo-opercular and default mode networks like rectus, anterior and middle gyrus, frontal superior gyrus, frontal superior medial gyrus, supplementary motor area, precuneus, cuneus, calcarine and occipital superior gyrus. The bilateral superior temporal gyrus, superior temporal pole and middle temporal pole play an inter-hemispheric modular connector role (see * in Figure 5). Five out of thirteen consistent connector hubs are located within inter-hemispheric modules. Interestingly, eight homologous brain areas are grouped together in either left (module 8) or right hemisphere (module 9). These areas are: hippocampus, parahippocampal gyrus, amygdala, inferior temporal gyrus, thalamus, pallidum, fusiform and lingual gyrus.

**Figure 5.**
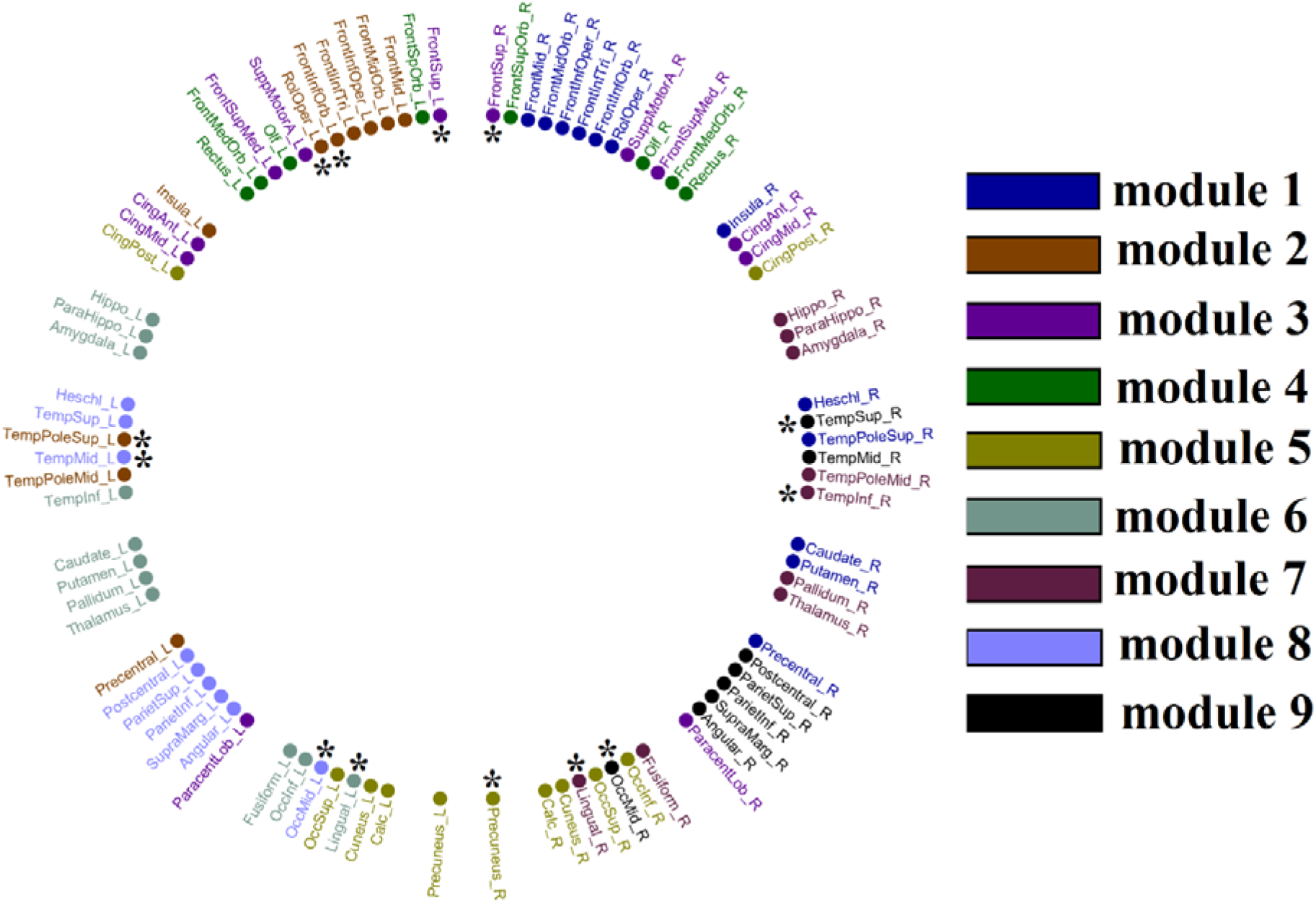
Topological Layout of Modular Assignment into the 90 AAL brain areas based on the community affiliation extracting from the consensus matrix related to 9-m OMST graph construction scheme and mscd_so community detection algorithm. With ‘*’, we denoted the connector hubs detected consistently across participants and repeat scans from the same combination of {mscd-so, 9-m OMST} (see section 3.5). This circular plot illustrates the 90 AAL brain areas into 45 of the left hemisphere on the left semi-circle and 45 of the right hemisphere on the right semi-circle. Our analysis gave nine communities/modules where each one is encoded with a different colour.

### 3.5 ICC of Nodal Participation Coefficient Index and Within-Module Z-score

Table 4 shows the group-averaged ICC of nodal Participation Coefficient *P*_i_. ICC values were first estimated per node and were then averaged across the 90 nodes for every pair of graph construction scheme and community detection algorithms. The highest ICC score was detected for the mscd_so algorithm combined with both the 9-m OMST and NS-thr graph construction schemes. On average across the seven graph construction schemes, the mscd_so algorithm also demonstrated the highest average ICC score. However, these findings did not reach statistical significance. Two-way ANOVA revealed no main effect of network-averaged of Participation Coefficient index across the four community detection algorithms (df =3, F = 2.86, p = 0.05) no main effect of the graph construction and no significant interaction term.

**Table 4.** ICC of Nodal Participation Coefficient index across every combination of graph construction scheme with community detection algorithms. We denote the top ranked values in bold letters.

|  | NS-OMST | NS+FA<br>OMST | 9-m<br>OMST | NS-thr | NS-t/FA-w | NS-t/MD-w | FA-t/NS-w |
| --- | --- | --- | --- | --- | --- | --- | --- |
| mscd_afg | 0.54 | 0.50 | 0.62 | **0.67** | 0.58 | 0.60 | 0.38 |
| mscd_rb | 0.61 | 0.55 | 0.68 | 0.73 | 0.67 | 0.65 | 0.46 |
| mscd_rn | 0.58 | 0.55 | 0.68 | 0.67 | 0.61 | 0.61 | 0.43 |
| mscd_so | 0.69 | 0.70 | **0.80** | **0.80** | **0.77** | **0.76** | 0.43 |

Figure 6 shows the nodal ICC for the {mscd_so,9-m OMST} and {mscd_afg, NS-thr} pairs. Applying a Wilcoxon Rank-Sum-test between the two sets of 90 ICCs, we detected a significantly higher ICC for the mscd_so – 9-m OMST pair (p-value = 0.041).

**Figure 6.**
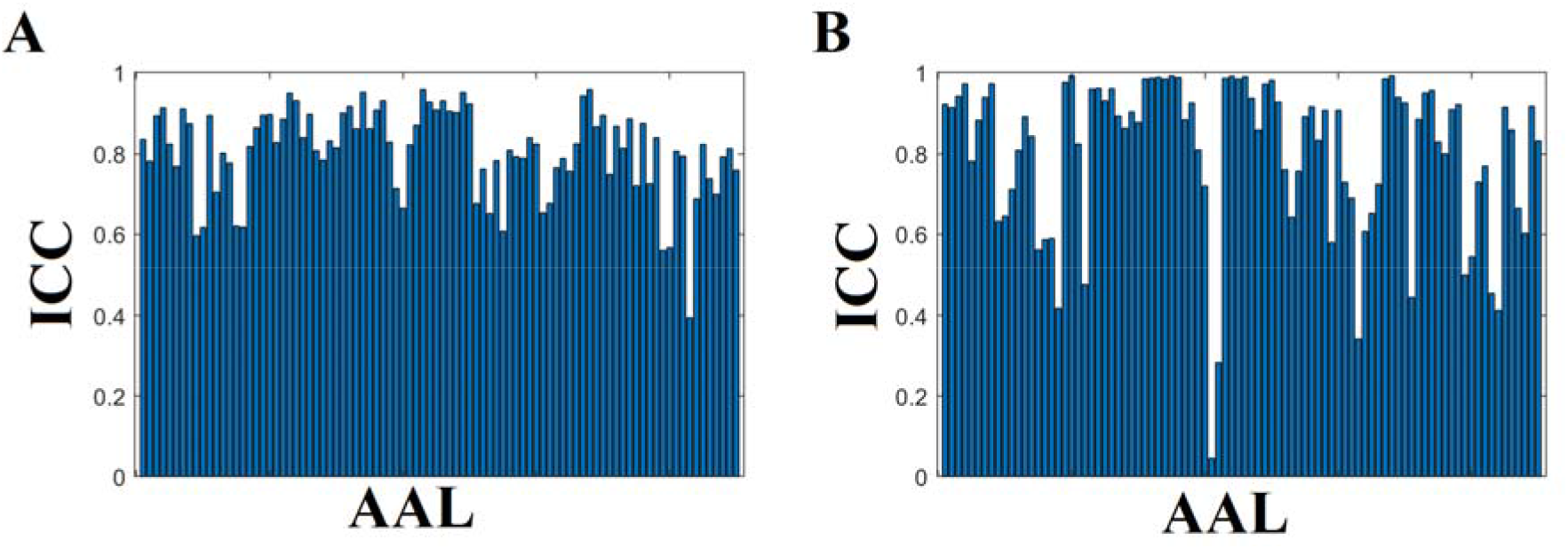
ICC of Nodal Participation Coefficient index for the best combinations of graph construction scheme and community detection algorithm. A. mscd_so – 9-m OMST B. mscd_afg – NS-thr

Table 5 shows the group-averaged ICC of nodal within-module Z-scores, *z*_i_. ICC values were first estimated per node and were then averaged across the 90 brain areas for every pair of graph construction scheme and community detection algorithms. The highest ICC scores were found for the mscd_rb and mscd_so mode algorithms combined with the NS-thr graph construction scheme. On average across the seven graph construction schemes, the mscd_so mode algorithm demonstrated the highest average ICC score. These findings reached statistical significance. Two-way ANOVA revealed a main effect of network-averaged z_i_ across the four community detection algorithms (df =3, F = 5.89, p = 0.0037, corrected for multiple comparisons) but no main effect of the graph construction scheme, nor an interaction term. The mscd_rb community detection algorithm produced the highest ICC.

**Table 5.** ICC of Nodal Within-Module Z-score across every combination of graph construction scheme with community detection algorithms. We denote the top ranked values in bold letters.

|  | NS-OMST | NS+FA OMST | 9-m OMST | NS-thr | NS-t/FA-w | NS-t/MD-w | FA-t/NS-w |
| --- | --- | --- | --- | --- | --- | --- | --- |
| mscd_afg | 0.71 | 0.66 | 0.65 | 0.79 | 0.69 | 0.69 | 0.69 |
| mscd_rb | **0.74** | 0.66 | 0.64 | **0.81** | 0.72 | 0.72 | 0.73 |
| mscd_rn | 0.56 | 0.57 | 0.60 | 0.68 | 0.55 | 0.57 | 0.61 |
| mscd_so | 0.73 | 0.59 | 0.73 | **0.75** | 0.60 | 0.62 | 0.52 |

Figure 7 shows the nodal ICC for the {mscd_rb, NS-thr} and {mscd_so, NS-thr}. Applying a Wilcoxon Rank-Sum-test between the two sets of 90 ICCs, we found the mscd_rb algorithm had significantly higher ICC (p-value = 0.0335 x 10^−9^).

**Figure 7.**
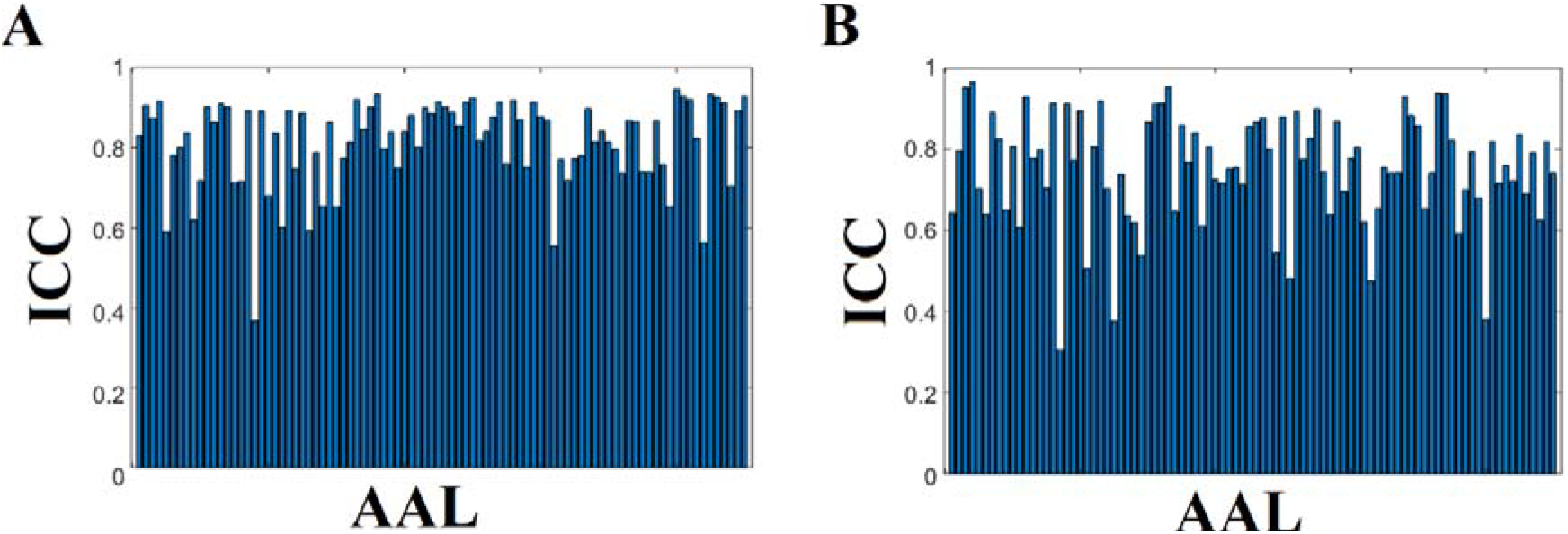
ICC of Nodal Within-Module Z-score for the best combinations of graph construction scheme and community detection algorithm. A. mscd_rb – NS-thr B. mscd_so – NS-thr

### 3.6 Reproducibility of Structural Hubs Detection based on Participation Coefficient Index and Within-Module Z-score

We estimated the Agreement Score of both connector and provincial hub detection across the cohort. The Agreement Score of connector hub detection across the cohort was higher than the Agreement Score for provincial hubs (Table 6 versus Table 7).

**Table 6.**
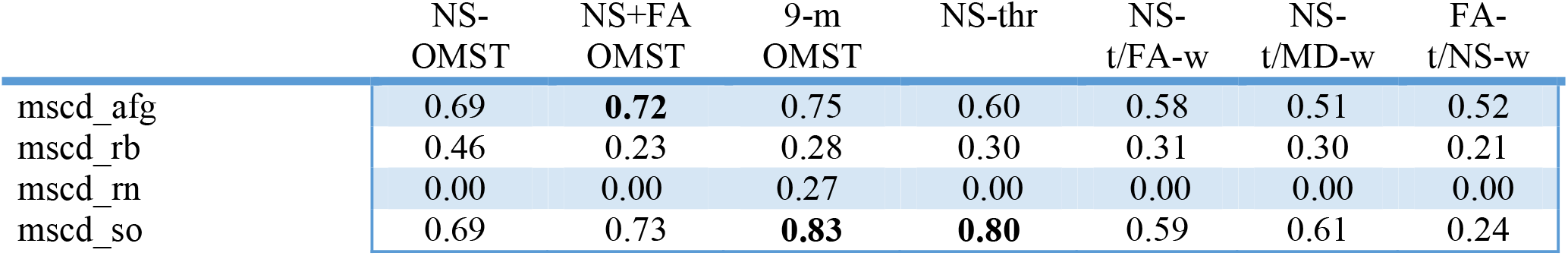
Agreement Scores of Provincial Hubs across every combination of graph construction scheme and community detection algorithm. We assigned with bold the top ranked values.

|  | NS-OMST | NS+FA<br>OMST | 9-m<br>OMST | NS-thr | NS-t/FA-w | NS-t/MD-w | FA-t/NS-w |
| --- | --- | --- | --- | --- | --- | --- | --- |
| mscd_afg | 0.69 | **0.72** | 0.75 | 0.60 | 0.58 | 0.51 | 0.52 |
| mscd_rb | 0.46 | 0.23 | 0.28 | 0.30 | 0.31 | 0.30 | 0.21 |
| mscd_rn | 0.00 | 0.00 | 0.27 | 0.00 | 0.00 | 0.00 | 0.00 |
| mscd_so | 0.69 | 0.73 | **0.83** | **0.80** | 0.59 | 0.61 | 0.24 |

**Table 7.** Agreement Scores of Connector Hubs across every combination of graph construction scheme and community detection algorithm. We assigned with bold the top ranked values.

|  | NS-OMST | NS+FA<br>OMST | 9-m<br>OMST | NS-thr | NS-t/FA-w | NS-t/MD-w | FA-t/NS-w |
| --- | --- | --- | --- | --- | --- | --- | --- |
| mscd_afg | 0.40 | 0.13 | 0.23 | 0.38 | 0.32 | 0.31 | 0.43 |
| mscd_rb | **0.65** | 0.42 | 0.45 | **0.67** | 0.57 | 0.58 | **0.80** |
| mscd_rn | 0.13 | 0.05 | 0.09 | 0.19 | 0.15 | 0.14 | 0.49 |
| mscd_so | 0.17 | 0.03 | 0.05 | 0.12 | 0.12 | 0.12 | 0.17 |

The highest Agreement Score for provincial hub detection was found for {mscd_so, 9-m OMST} (Table 6). On average across the seven graph construction schemes, the mscd_so algorithm demonstrated the highest average Agreement Score for provincial hub detection. Two-way ANOVA revealed a main effect of network-averaged Within-Module Z-score across the four community detection algorithms (df =3, F = 35.55, p = 0.542 x 10^−6^, corrected for multiple comparison) but no main effect of the graph construction schemes nor an interaction. The mscd_so community detection algorithm produced the highest Agreement Score for provincial hub detection.

The highest Agreement Score for connector hubs detection was found for {mscd_rb, FA-t/NS-w} (Table 7). On average across the seven graph construction schemes, the mscd_rb algorithm demonstrated the highest average Agreement Score of connector hub detection. These findings are supported statistically. Two-way ANOVA revealed a main effect of network-averaged Within-Module Z-score across the four community detection algorithms (df =3, F = 24.3, p = 0.188 x 10^−4^) but no effect of the graph construction schemes nor their interaction. The mscd_rb community detection algorithm produced the highest agreement score for connector hub detection.

Figure 8 shows the Agreement Score for connector hubs for the pair {mscd_rb, FA-t/NS-w}. The group of connector hubs is indicated alongside modular representation of consensus modules illustrated in Figure 5 and also in Table 8. Interestingly, five out of thirteen consistent connector hubs are located within the inter-hemispheric modules (see Figure 5). Our conclusion is that the combination of modular network metrics P_i_ and z_i_ succeeded in uncovering a consistent core of connector hubs but failed to detect provincial hubs consistently.

**Table 8.**
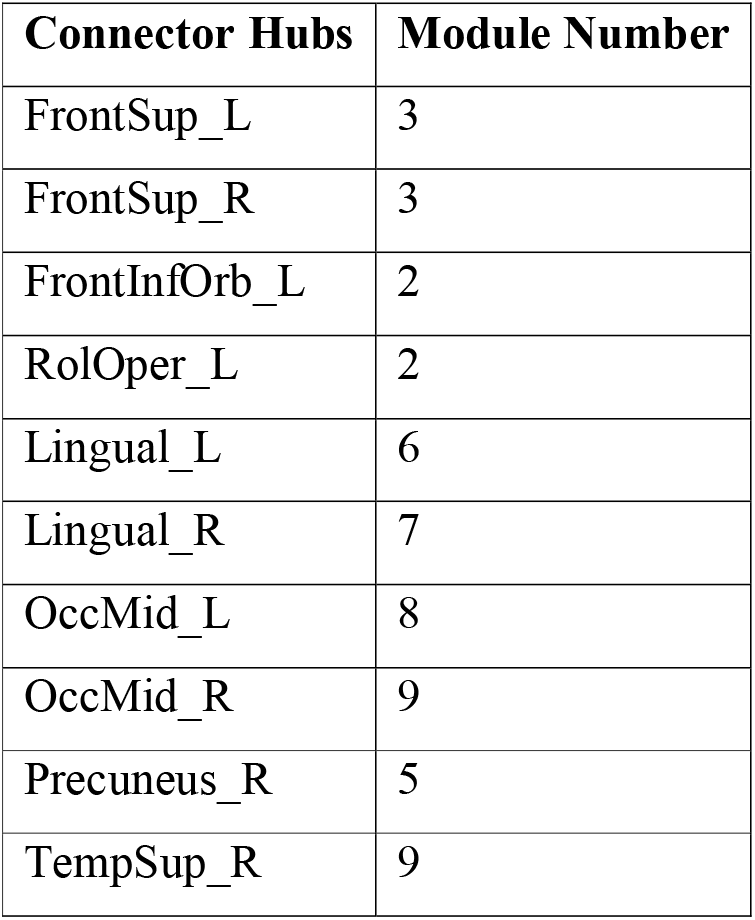
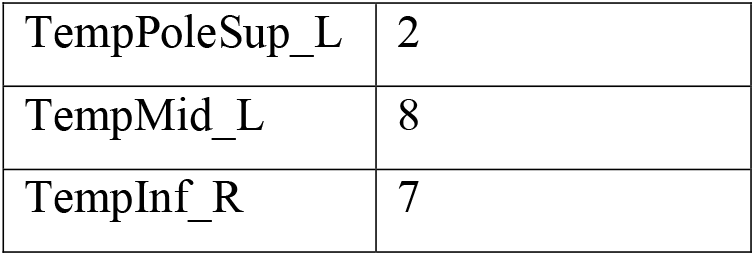
Consistent connector hubs aligned with the detected module number illustrated in Figure 5.

| Connector Hubs | Module Number |
| --- | --- |
| FrontSup_L | 3 |
| FrontSup_R | 3 |
| FrontInfOrb_L | 2 |
| RolOper_L | 2 |
| Lingual_L | 6 |
| Lingual_R | 7 |
| OccMid_L | 8 |
| OccMid_R | 9 |
| Precuneus_R | 5 |
| TempSup_R | 9 |
| TempPoleSup_L | 2 |
| TempMid_L | 8 |
| TempInf_R | 7 |

**Figure 8.**
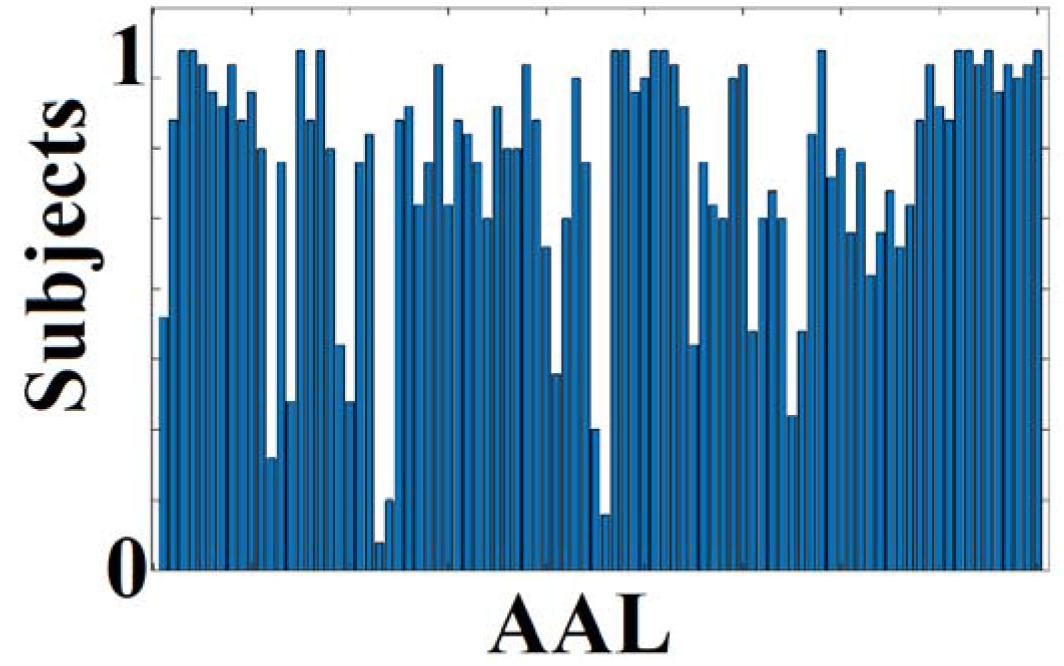
Agreement Score of Connector Hubs for the best pair of {mscd_rb,FA-t/NS-w}.

## 4. Discussion

We have presented the first extensive study in the literature on the robustness of community detection in structural brain networks by exploring different graph-construction schemes (previously shown to exhibit high repeatability themselves) and various community detection algorithms. Our main findings have direct implications for longitudinal studies and studies comparing healthy controls versus diseased populations.

The key findings of our analysis can be summarized as follows:

1. The repeatability of community affiliations depends heavily on the combination of graph-construction scheme and community detection algorithm. All previously reported studies of network communities adopted a specific pair of graph-construction scheme and community detection algorithm, with the majority of them focused on Newman’s modularity objective criterion (Newman, 2006; Sporns and Betzel, 2016; Betzel et al., 2017). Based on our first criterion of high repeatability of community affiliation between the two scans and across the cohort (NMI > 0.9) supported statistically via bootstrapping (p = 0.0001), we identified four community detection algorithms as the best choices:

A. mscd_agb
B. mscd_rb
C. mscd_rn
D. mscd_so These four algorithms gave excellent repeatability across the entire set of graph-construction schemes (see Table 2 and Figure 4).
2. Complementary to the repeatability of the community partitions, we assessed their quality (second criterion). The majority of the graph partition algorithms used here (28 out of 33) estimated a quality index in combination with the community partition. By comparing the quality index of the original community partition to that derived from 1000 surrogate null graph versions, we computed their significance values to be in the range 0.013 < P < 0.021. For the soft clustering algorithms, where this quality index was absent, we estimated the mean clustering distance (mean NMI) between the original partition and partitions derived from 1000 surrogate null graph versions of the original graph. The mean NMI was below 0.1. Our findings indicated community detection at the mesoscale is repeatable and, as the Q quality for the four selected algorithms reached statistically significant levels, is of high quality.
3. Our third criterion was the high similarity of consensus community affiliation with individual community affiliations. The majority of the neuroimaging studies that employed a single graph-construction scheme and one community detection algorithm did not evaluate their findings under this framework (Ryali et al., 2015; Rosero et al., 2017; Akiki and Abdallah, 2019). The consensus matrix integrates community affiliations across the entire cohort counting the number of times two nodes are assigned to the same community. To reveal a representative community affiliation, we have to apply a community detection algorithm. However, almost all previous studies report this representative community affiliation was determined without estimating its similarity with *individual* community affiliations. Following a two-way ANOVA, we observed an effect of group-averaged similarity of individual community partitions with consensus community partition across the four community detection algorithms. Of the four community-detection algorithms, the mscd_so community detection algorithm produced the highest group-averaged similarity of individual community partitions with consensus community partition. We found the highest similarity between individual community partitions and representative community partition for the {mscd_so, 9-m OMST} pair. The second highest similarity was detected for {mscd_afg, 9-m OMST} pair.
4. Two-way ANOVA reported an effect of group-averaged similarity of individual community partitions with consensus community partition across the four community detection algorithms (df =3, F = 54.14, p = 0.783 x 10^−9^, corrected for multiple comparisons) but there was no effect of the graph construction scheme or their interaction. The mean group-averaged similarity across the seven graph construction schemes allowed us to rank the four graph construction schemes.
5. An important result of our analysis is that soft clustering community detection algorithms gave the least repeatable results. Therefore, we recommend the use of hard-clustering algorithms for the detection of brain communities, at least when using the AAL template.
6. The best combination of graph-construction scheme (9-m OMST) and community detection algorithm (mscd_so) revealed nine distinct modules (as illustrated topologically in Figure 5) with interesting findings:

A. Modules 1, 7 and 9 group together brain areas located exclusively within the left hemisphere while modules 2, 6 and 8 group together brain areas located exclusively within the right hemisphere. Modules 3, 4 and 5 group brain areas from both hemispheres together.
B. Modules 3 – 5 that integrate brain areas from both hemispheres involve bilateral ROIs from fronto-parietal, cingulo-opercular and default mode network like rectus, anterior and middle gyrus, frontal superior gyrus, frontal superior medial gyrus, supplementary motor area, precuneus, cuneus, calcarine and occipital superior gyrus. A multi-tasking fMRI study has previously suggested the flexible role of the fronto-parietal network in cognitive control and adaptive demands of cognitive tasks (Cole et al., 2013).
C. Five out of thirteen consistent connector hubs are located within inter-hemispheric modules 3 – 5 supporting their inter-connecting role (Figure 5).
D. Interestingly, eight homologous brain areas were grouped together in either left (module 8) or right hemisphere (module 9). These areas are: hippocampus, parahippocampal gyrus, amygdala, inferior temporal gyrus, thalamus, pallidum, fusiform and lingual. Lesions of hippocampus, parahippocampal gyrus, amygdala and fusiform gyrus in participants with temporal lobe epilepsy caused an impaired associative memory in learning tasks that require learning and recall of objects and faces (Weniger et al., 2004). These four brain areas including the thalamus are those most consistently implicated in neurodegenerative dementias, especially in Alzheimer’s Disease, even at an early stage (Manuello et al., 2018).
E. The bilateral superior temporal gyrus, superior temporal pole and middle temporal pole play an inter-hemispheric integration role. Inter-hemispheric functional connections between temporal lobes predict language impairment in adolescents born preterm (Northam et al., 2012). Phonological awareness, a key factor in reading acquisition was positively correlated with radial diffusivity of the interhemispheric pathways connecting temporal lobes (Dougherty et al. 2007). This bilateral temporal module could play a key role in many functions and dysfunctions.
7. The core of our study was an extensive analysis to identify the optimal pair of graph-construction scheme and community detection algorithm. The choice of this pairing will also affect repeatability of connector and provincial hub detection based on the participation coefficient score P_i_ and the within-module z-score z_i_. Our results revealed a high repeatability of nodal P_i_ with the mscd_so algorithm across the seven graph construction schemes. The highest ICC score was reached for the {mscd_so, 9-m OMST} pair. A significantly higher repeatability of nodal z_i_ was found for mscd_so algorithm compared to the rest of the community detection algorithms. The highest ICC was achieved for the pairs {mscd_rb, NS-thr} and {mscd_so, NS-thr}.
8. The Agreement Score was high only for connector and not for provincial hubs using both modular network metrics. The Agreement Score was statistically higher for the {mscd_so, 9-m OMST} pair than for the rest of the community detection algorithms. We detected a group of thirteen repeatable connector hubs across the cohort, but no provincial hubs. Based on our results, we therefore recommend to not use these modular network metrics for the detection of provincial hubs, at least when using the AAL atlas. The designation of a brain node as a hub depends also on the scale at which brain networks are constructed. Many brain areas in a basic atlas template group together functionally heterogenous subareas and it is possible that a finer-grained parcellation may affect the nodes’ classification as a hub or not. For example, the thalamus, despite comprising 50 – 60 specialized sub-nuclei (Herrero et al., 2002) is in many studies, including ours, treated as a single node.

In our previous study on the same cohort, we focused on the repeatability of network topologies focusing on edge weights and graph theoretical metrics. We demonstrated that network topology and edge weights are repeatable, but the repeatability depends on the graph-construction scheme (Messaritaki et al., 2019b). The important finding in this work is that the repeatability of network topologies and edge weights does not guarantee the repeatability of community detection at the mesoscale. In the present study, we focused on this important tool for mesoscale network topological investigations, and the detection of robust communities in structural brain networks over the same participants. To the best of our knowledge, this is the first study in the literature that explores the robustness of community detection over a large set of graph-construction schemes (seven) and community detection algorithms (thirty-three). Our analysis detected an optimal pair of {mscd_so, 9-m OMST} that fulfills the three basic criteria: high repeatability of community affiliations between the two scan-sessions, quality over surrogate null graph partitions and high similarity of group community affiliations with the individual community affiliations. To the best of our knowledge, this is the first time that the second and third criterion were used for the validation of representative consensus community affiliation (this includes studies using a single graph-construction scheme and community detection algorithm).

Running the comparison study for the whole set of thirty-three graph partition algorithms (including graph partition of the original graph and 1000 surrogate null models) takes a few hours on a personal computer. We suggest to the neuroscience community to always run such an analysis over an in-house test-retest dataset acquired with the same settings as in the targeted dataset. Optimizing the set of algorithms over the test-retest study will increase the chance of repeatability of findings over the single-scan dataset. This process will increase the reproducibility of research findings, especially important for cross-sectional studies.

Our study has a few limitations. This dataset involves a specific data acquisition protocol and a specific tractography algorithm. We recommend following our analysis for every study because such an investigation could improve the repeatability and reproducibility of the findings at the mesoscale while also increasing the power of the study at the nodal and network level (Messaritaki et al., 2019b). Additionally, in our study we used only one of the three available b-values to perform the tractography. This was mainly done in order to supplement our previous work (Messaritaki et al., 2019), and we chose the b-value of 2000 s/mm^2^, because this value provides a balance between the b-value being sufficiently high to resolve crossing fibers with CSD, while at the same time being ensuring sufficient SNR in the signal for robust measurements, and that higher-order effects of the diffusion do not need to be taken into account when calculating the diffusion metrics. Using one b-value also reflects acquisition protocols routinely used in other studies, and therefore makes our work more applicable to the general literature. At the same time, tractography results could be improved by combining data from all avaialble b-values, and the implications of using different community detection algorithms in those cases should be explored as well. Moreover, three or more scan sessions would be also desirable to get a more robust assessment of repeatability. Scanning the same participant on different scanners and /or with different protocols would also allow assessment of reproducibility as well as repeatability. Lastly, the reproducibility of estimates of structural brain networks is affected by the resolution of the MR data (Vaessen et al., 2010), the parcellation scheme used (Bassett et al., 2011), the interval time between the scan sessions and others.

## 5. Conclusions

In this study, we compared several graph-construction schemes and community detection algorithms for the detection of reproducible communities in structural brain networks. Our extensive analysis showed that every choice in both groups of algorithms exhibits different reproducibility in community detection algorithms, as well as in connector/provincial hubs detection. Our analysis indicates that our analytic pathway should be adopted and performed in every study in order to extract reliable results at the mesoscale of structural brain networks.

## Author Contributions

Conceptualization: SID

Methodology:SID

Software:SID

Validation:SID

Formal analysis:SID, EM

Formal Analysis: SID

Investigation:SID

Resources: The original diffusion MRI are free available from the Human Connectome project. The analysis of tractography and the construction of structural brain networks has been realized by EM.

Data curation:EM,SID

Roles/Writing - original draft: SID,

Writing - review & editing: EM,DJ

Funding acquisition:EM,SID

## Data and code availability

The HCP test-retest data is freely available as listed above.

The code used to generate the graphs for the structural brain networks with the OMST schemes is available at: https://github.com/stdimitr/multi-group-analysis-OMST-GDD. The structural brain networks and the code used to perform the reproducibility analysis will be released as soon as the paper will be accepted from author’s github website https://github.com/stdimitr.

Source code of community detection algorithms are provided on the Dr.Le Martelot’s personal homepage, author’s homepage and also were implemented by our team. The collection of the whole set of the algorithms will be reported in our github homepage.

## Acknowledgements

We are grateful to the Human Connectome Project for making the test-retest data freely available.

“Data were provided [in part] by the Human Connectome Project, WU-Minn Consortium (Principal Investigators: David Van Essen and Kamil Ugurbil; 1U54MH091657) funded by the 16 NIH Institutes and Centers that support the NIH Blueprint for Neuroscience Research; and by the McDonnell Center for Systems Neuroscience at Washington University.”

The work was partly funded under the BRAIN Biomedical Research Unit, which is funded by the Welsh Government through Health and Care Research Wales. DKJ is supported by a Wellcome Trust Investigator Award (096646/Z/11/Z) and a Wellcome Trust Strategic Award (<u>104943/Z/14/Z</u>). SID was supported by MRC grant <u>MR/K004360/1</u> and a Marie Sklodowska-Curie COFUND EU-UK Research Fellowship. EM is supported by a Wellcome Trust ISSF Research Fellowship at Cardiff University (204824/Z/16/Z).

## Appendix 1. Detecting Structural Hubs based on Nodal Participation Coefficient P_i_ and Within-Module Z-Score z_i_

By recovering the community partition and estimating the participation coefficient, we can classify brain hubs into provincial and connector hubs (Guimera and Amaral,2005). ‘Provincial hubs’ are high-degree nodes that primarily connect to nodes in the same module. ‘Connector hubs’ are high-degree nodes that show a diverse connectivity profile by connecting to several different modules within the network. Brain hubs are important brain areas that are vulnerable and susceptible to disconnection and dysfunction in brain disorders. Rich club organization of structural hubs supports the robustness of inter-hub connections and promotes the efficient information exchange between brain areas and its integration across the brain (van den Heuvel and Sporns,2011).

The distinction of nodes into hubs and non-hubs by a combination of network topology and community affiliation is supported by a pair of network metrics called: participation coefficient *P_i_* and within-module z-score, *z_i_*. This definition has been first reported by Guimera and Amaral (2005). Here, we first reported the reliability of these nodal metrics in structural brain networks.

The degree of a node *i* is defined as, where *A_ij_* is the adjacency matrix of the graph. Within-module z-score for node *i* is defined as

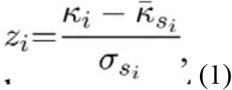

where κi is the number of edges of node *i* to other nodes in its module *s*_i_ *s_i_* is the average of κ over all the nodes in *s_i_*, and σ*si* is the standard deviation of κ in *s_i_*.

The participation coefficient P_i_, for node *i* is defined as

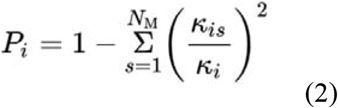

where *k_is_* is the number of links of node *i* to nodes in module *s*, and *k_i_* is the total degree of node *i*. The participation coefficient of a node is therefore close to 1 if its links are uniformly distributed among all the modules and zero if all its links are within its own module.

## Provincial Hubs

If a node with a large degree, *k* » 1, has at least 5*/*6 of its links within the module, then it follows that *P* = 1 − (5*/*6)^2^ − (*k/*6)(1*/k*^2^) = 0.31 − 1*/*(6*k*) ≈ 0.30. i.e., *P < 0.3*

## Connector Hubs

If a node with a large degree has at least half of its links within the module, then it follows that *P* = 1 − 1*/*4 − (*k/*2)(1*/k*^2^) = 0.75 − 1*/*(2*k*). Since *k* » 1, *P* < 0.75 for such nodes.

Both provincial and connector hubs demonstrate a high within-module z-score which means that they have many within-module edges. In this work, we used the threshold originally proposed by Guimera and Amaral, (2005) for the z_i_ dimension as z_i_ > 2.5 for both types of studied hubs (see Figure 5 in Guimera and Amaral, (2005)). In the *P_i_* dimension, we defined a node as provincial hub if *P_i_* <= 0.3 and as connector hub if 0.3 < *P_i_* < 0.75.

The intra-class correlation coefficient (ICC) was estimated for every nodal participation coefficient index, *P_i_,* and within-module z-score z_i_ across the cohort and for every selected pair of graph construction scheme and community detection algorithm that showed higher group-averaged community similarity (NMI > 0.9).

## Appendix 2. Graph partition Algorithms

We described briefly the thirty-three graph-partition algorithms used in the present study.

Hard clustering algorithms are divided into three groups:

## A. Fast multi-scale community detection algorithms

1. (<u>fast_mo</u>): Fast greedy modularity optimisation algorithm based on the multi-scale algorithm but optimised for modularity (mono-scale) (Le Martelot and Hankin, 2012a,b).
2. (<u>mscd_afg</u>): Fast multi-scale community detection algorithm using the criterion from Arenas et al. (2008)
3. (<u>mscd_hslsw</u>): Fast multi-scale community detection algorithm using the criterion from Huang et al. (2011)
4. (<u>mscd_lfk</u>): Fast multi-scale community detection algorithm using the criterion from Lancichinetti et al. (2009b)
5. (<u>mscd_rb</u>): Fast multi-scale community detection algorithm using the criterion from Reichardt & Bornholdt, (2006)
6. (<u>mscd_rn</u>): Fast multi-scale community detection algorithm using the criterion from Ronhovde & Nussinov (2009)
7. (<u>mscd_so</u>): Fast multi-scale community detection algorithm using stability optimisation (Le Martelot and Hankin, 2012).

## B. Multi-Scale Community Detection algorithms using Stability as Optimisation Criterion in a Greedy Algorithm and Multi-Scale Community Detection using Stability Optimisation as described in (Le Martelot and Hankin,2011)

8. (<u>gso_discrete</u>): Greedy stability optimisation using the Markov chain model.
9. (<u>gso_continuous</u>): Greedy stability optimisation using a time-continuous Markov process.
10. (<u>msgso_discrete</u>): Multi-step greedy stability optimisation using the Markov chain model.
11. (<u>msgso_continuous</u>): Multi-step greedy stability optimisation using a time-continuous Markov process.
12. (<u>rgso_discrete</u>): Randomised greedy stability optimisation using the Markov chain model.
13. (<u>rgso_continuous</u>): Randomised greedy stability optimisation using a time-continuous Markov process.
14. (<u>gso_discrete_markovian</u>): Front-end function calling any of the aforementioned algorithms based on discrete Markovian model. We specified a pre-processing phase removing nodes with a single neighbour and a post-processing phased based on the Kernighan-Lin algorithm adapted for stability (Kernighan and Lin,1970).
15. (<u>gso_discrete_continuous</u>): Same as above but based on continuous Markovian model.
16. (<u>Newman</u>): Greedy modularity optimisation with Newman’s fast algorithm.
17. (<u>Danon</u>): Greedy optimisation of Danon et al.’s criterion (Danon et al., 2006).
18. (<u>Louvain</u>): Uses the Louvain method (Blondel et al.,2008)
19. (<u>Louvain_modularity</u>): Uses the Louvain method to optimise modularity criterion with gamma=0.5.
20. (<u>Louvain_so</u>): Uses the Louvain method to optimise ‘stability’ criterion with Markovian time
21. (<u>reichardt</u>): Greedy optimisation of Reichardt & Bornholdt’s criterion (Reichardt and BornHoldt, 2006).
22. (<u>ronhovde</u>): Greedy optimisation of Ronhovde & Nussinov’s criterion (Reichardt and BornHoldt, 2006)

## C. Hard community detection algorithms involving state-of-the-art graph partition algorithms

23. (<u>shi_malik</u>): From tens of available spectral clustering algorithms, we adopted the algorithm from Shi and Malik, (2000)
24. (<u>dominant_sets</u>): Dominant sets (Pavan and Pelillo, 2007). We have adopted this algorithm in our previous studies (Dimitriadis et al., 2009,2012)
25. (<u>modularity</u>): The algorithm by Newman, 2006
26. (<u>infomap</u>): An important algorithm for identifying communities in large graph-oriented problems and synthetic graphs (Rosvall M, Bergstrom CT, 2008))

## The soft clustering algorithms we considered are

27. (<u>ogso_discrete</u>): Greedy stabilisation optimisation with overlapping communities for discrete Markov Model. The network is first converted to an edge-graph and then community detection is performed on the edge-graph
28. (<u>ogso_continuous</u>): Greedy stabilisation optimisation with overlapping communities for continuous Markov Model. The network is first converted to an edge-graph and then community detection is performed on the edge-graph
29. (<u>link</u>): Link Communities: The network is first converted to an edge-graph and then community detection is performed on the edge-graph. In an edge-graph, nodes refer to edges of the original SBN while edges denote edges between pair of connected edges of the original SBN. This algorithm uncovers overlapping community structure via hierarchical clustering of network links (Ahn et al., 2010). We used single-linkage criterion.
30. (<u>link_complete</u>): Link Communities: Same as above but we used complete-linkage criterion.
31. (<u>nnmf</u>): Non-negative Matrix Factorization (NNMF): (Psorakis et al., 2011)
32. (<u>k-cliques</u>): This algorithm partitions a graph into soft communities using the simplest subgraph forms, the k-cliques. Here, we used 3-cliques (Palla et al., 2005)
33. (<u>orthogonal nonnegative matrix t-factorizations</u>): Orthogonal nonnegative matrix t-factorizations (Ding et al., 2006)

For the first set of multiscale community detection algorithms, we considered the following set of resolution parameter ps = [0.1 0.15, …, 0.55, 0.60]. For the discrete and continuous Markovian time series, we searched over ts = [0.1, 0.02, …, 0.9,1.0]. Finally, for Louvain’s modularity method and Reichardt’s method, we considered γ = [0.05, 0.1, 0.15, …,0.95,1.5]. Τhe aforementioned parameters were optimized over the highest group-averaged community affiliation similarity between the two scans estimated with NMI. For further details, see the following section.

For the Louvain’s methods, we ran the algorithms 1000 times and we followed the construction of consensus matrix approach. We described this approach in section 2.3.8 for the construction of consensus matrix across participants and scans.

## Appendix 3. Normalized Mutual Information: Graph – partition similarity

To assess the reproducibility of the thirty-three community detection techniques across the seven graph-construction schemes and repeat-scan sessions, we first quantified the similarity between the community partitions from the two scan sessions separately for every participant using the Normalized Mutual Information (NMI) (Alexander-Bloch et al. 2012), defined as follows (Lancichinetti and Fortunato,2009a):

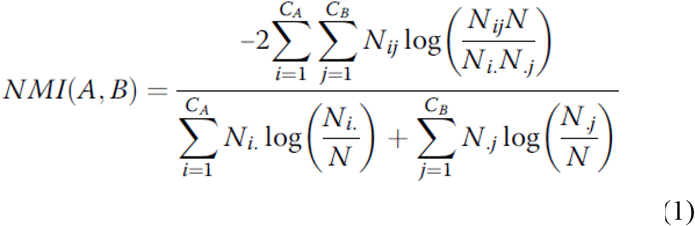

where *A* and *B* are the community partitions of two SBNs from the two scan sessions while *C_A_, C_B_* are the number of communities in partition *A* and *B*, correspondingly. *N* denotes the number of nodes (here 90), while *N_ij_* is the overlap between *A*’s and *B*’s communities *i* and *j* which practically means the number of common nodes between the two partitions. *N_i_* and *N_j_* are the total number of nodes in *A*’s and *B’s* communities *i* and *j* respectively. The NMI ranges from 0 to 1 where 0 corresponds to two independent partitions and 1 to identical partitions. This definition was used for hard community partition comparisons while for soft community partition, we adopted the homologue definition of NMI tailored to soft graph clustering (Lancichinetti et al., 2009a).

We calculated NMI values between every possible pair of scans and for each of the seven graph-construction schemes and the thirty-three community detection algorithms giving an space of {7 graph-construction schemes x 33 community detection algorithms}. The NMI was then averaged across the 37 participants, to create a group-averaged NMI, and ranked the community detection algorithms with high test-retest reproducibility (group-averaged values NMI > 0.9) in at least one of the seven graph construction schemes (see Figure 4).

To quantify the statistical significance of group-mean community similarity quantified with NMI, we adopted a nonparametric test via bootstrapping procedure (Dimitriadis et al., 2012). Practically, the outcome of every community detection algorithm in our cohort is a matrix (dimensions {(no of participants) × (no of nodes)}) with elements containing integer numbers assigned to every detected community. Analysing the scan-rescan data gives two such matrices, one per scan session. The bootstrapping procedure comprises:

a. The two matrices (dimensions {(no of participants) × (no of nodes)}) tabulating the community affiliations across the cohort are integrated into a single matrix (dimensions {2 x no of participants x no of nodes})
b. The rows of this integrated matrix are shuffled, mixing the community affiliations of participants across scan sessions
c. The integrated matrix is divided into two matrices of equal size {no of participants x no of nodes} and the group mean NMI between those two shuffled matrices is estimated.
d. Steps (b) and (c) are repeated 10,000 times
e. A P-value is assigned to every graph construction scheme - community detection algorithm pair by counting the number of times the permuted between-scan group mean similarity exceeds the actual between-scan group mean similarity, divided by the number of permutations (here 10,000)

